# Comprehensive Genotoxicity and Subchronic Oral Toxicity Evaluation of Kumamolisin from Bacillus sp. MN-32 for Human Use

**DOI:** 10.64898/2025.12.25.696157

**Authors:** Wai Shun Mak, V Pradeep, G Deshmukh, MC Umesh, GP Chethankumara, SJ Dev, T Rangappa, BM Shuheb, CM Doreswamy, SB Venkataramaiah, S Seekallu

## Abstract

Kumamolisin from Bacillus sp. MN-32 (produced by E. coli B/BL21 background) is an acid-active sedolisin protease developed for food uses; its systemic safety was assessed using GLP (Good-laboratory-practice) *in vitro* genotoxicity assays and repeated-dose oral studies. A bacterial reverse-mutation assay (OECD TG 471; Salmonella TA98, TA100, TA102, TA1535, TA1537; plate-incorporation and preincubation; ±S9) showed no concentration-related increases in revertants up to 5000 µg total organic solids (TOS)/plate. A mammalian cell micronucleus test (OECD TG 487; human lymphocytes; 4 h ±S9 and 24 h −S9) showed no significant increases in micronucleated binucleated cells at 1250–5000 µg TOS/mL, with cytotoxicity ≤ 21.6% by CBPI (Cytokinesis-Block Proliferation Index). In 28-day and 90-day repeated dose toxicity studies Sprague Dawley rat studies (OECD TG 407/408; n = 10/sex/group) at 0, 500, 1000, and 2000 mg TOS/kg bw/day by gavage (10 mL/kg/day), survival was 100% and in-life observations, ophthalmology, neurobehavior, urinalysis, hematology, coagulation, and clinical chemistry were unremarkable; isolated statistical differences (e.g., male total protein, BUN, glucose, LDL; calcium; female phosphorus) were marginal, within historical ranges, non-monotonic, and lacked histopathology correlates. Organ-weight differences were single-sex and non-dose-dependent. No test-item-related lesions were detected. The NOAEL was 2000 mg TOS/kg bw/day ((highest dose tested) in both the 28-day and 90-day studies, supporting oral safety for food-use exposure.

## Introduction

Proteases are core processing tools across food applications. They hydrolyze proteins to modify texture, improve digestibility, generate flavor peptides, debitter plant proteins, condition dough, tenderize meat, and prevent protein haze in beverages. Industrial workflows deploy acidic, neutral, and alkaline proteases to match unit-operation pH and temperature profiles in bakery, dairy, meat, and beverage lines, leveraging fermentation for high-purity preparations and consistent activity (1).

Kumamolisin from Bacillus sp. MN-32 is an acid-active, thermotolerant endopeptidase classified in the sedolisin family (MEROPS S53; EC 3.4.21.123) (2–4). Sedolisins are serine–carboxyl peptidases that operate with a catalytic Ser–Glu–Asp (S–E–D) triad and exhibit optima at low pH; they are pepstatin-insensitive and mechanistically distinct from aspartic proteases, which use catalytic Asp dyads (2). The MN-32 enzyme was originally described as a thermostable, pepstatin-insensitive acid proteinase produced by a Bacillus strain, consistent with sedolisin biochemistry and application in acidic matrices (5).

Kumamolisin is synthesized as a pro-enzyme comprising inhibitory propeptide and catalytic domain. The prodomain occludes the active site and functions as an intramolecular chaperone; autocatalytic processing under acidic conditions yields the mature enzyme (2,6). Structural studies resolved both the zymogen (pro-kumamolisin) and mature enzyme from Bacillus sp. MN-32. Crystallography shows a monomeric catalytic domain with an apparent mass near 40 kDa for the mature enzyme, while the zymogen is larger (≈55 kDa), reflecting the presence of the prodomain (6–8). Structures capture the prodomain in the active-site cleft and identify bound ions consistent with enhanced stability at low pH (6–8).

These biochemical and structural features make MN-32 kumamolisin a practical candidate for unit operations that require acid stability and thermal tolerance. Examples include dough softening and extensibility control, accelerated proteolysis in dairy fermentations, flavor-peptide generation from plant or dairy proteins, meat tenderization, and colloidal stabilization in beverages by targeted haze-forming protein cleavage—use cases where brief low-pH or heat exposures and broad substrate scope are advantageous (1–2).

To characterize systemic hazards relevant to oral exposure from food uses, we evaluated the toxicology profile of Bacillus sp. MN-32 kumamolisin (preparation produced in *E. coli* B/BL21 background) using guideline studies: a bacterial reverse-mutation assay and an *in vitro* mammalian cell micronucleus test for genotoxicity, and 28-day and 90-day repeated-dose oral toxicity studies in rats to derive a no-observed-adverse-effect level (NOAEL). Study designs followed OECD Test Guidelines for the micronucleus assay, Ames test, 28-day and the 90-day rodent study, enabling quantitative risk characterization (9–11).

## Materials and Methods

### Test article and vehicle

The test article was a kumamolisin preparation from Bacillus sp. MN32 produced in *E. coli* B/BL21-DE3, supplied as a white to pale yellow hygroscopic solid (purity 86% total protein by HPLC 280 nm; total organic solids, TOS, 58%). Batch T/CP24S3/U2/24/001/001. Protease activity (hemoglobin-based) was 7586 U/mg TOS where %TOS = 100 – (A+W+D), where A = ash, W = % water, and D = % diluent and carrier. The lot pH was 6.4 and the material was water soluble. The article was stored at 22 ± 3 °C. All concentrations and doses are expressed as mg TOS. Ultrapure water (type 1) served as vehicle.

### Dose formulation, stability, and homogeneity (28- and 90-day studies)

For the 28- and 90-day oral toxicity studies, dose formulations of kumamolisin were prepared at 50, 100, and 200 mg TOS/mL to deliver 500, 1000, and 2000 mg TOS/kg bw/day at 10 mL/kg. A TOS correction factor of 1.724 was applied when weighing the bulk powder. The weighed solid was triturated with vehicle to a paste, brought to volume with ultrapure water, vortexed, and sonicated for 10 min. Formulations were stored at 2–8 °C and used within validated stability windows (7 h at 22 ± 3 °C; 72 h at 2–8 °C). Final pH and appearance on first preparation were recorded: vehicle, pH 7.58 (colorless); 50 mg TOS/mL, pH 6.24 (off-white suspension); 100 mg TOS/mL, pH 6.10 (off-white suspension); 200 mg TOS/mL, pH 6.02 (off-white suspension). Homogeneity and dose confirmation (top/middle/bottom sampling; ≈1 mL per layer) were assessed using a validated HPLC-PDA method for total protein. Acceptance criteria were 100 ± 30% mean recovery and ≤20% RSD, with vehicle blanks required to be negative. For the 28-day study, mean recoveries (±SD; RSD) at Day 0 were 102.50 ± 0.38% (0.37%) for 50 mg/mL, 102.38 ± 1.11% (1.08%) for 100 mg/mL, and 97.75 ± 0.81% (0.83%) for 200 mg/mL; Week 4 values were 96.24 ± 1.15% (1.19%), 99.40 ± 0.80% (0.80%), and 97.62 ± 0.96% (0.99%), respectively. For the 90-day study, Day 0 recoveries were 102.50 ± 0.38% (0.37%), 102.38 ± 1.11% (1.08%), and 97.75 ± 0.81% (0.83%) for 50, 100, and 200 mg/mL, and Day 90 recoveries were 101.62 ± 0.74% (0.73%), 100.95 ± 0.98% (0.97%), and 99.43 ± 0.90% (0.91%), respectively.

### In vitro genotoxicity

Bacterial reverse-mutation assay (Ames; OECD TG 471)

Mutagenicity was assessed per OECD Test No. 471 using Salmonella typhimurium TA98, TA100, TA102, TA1535, and TA1537 (11). Plate incorporation and preincubation methods were run with and without rat liver S9 (10% v/v). Test concentrations were 15.8, 50, 158, 500, 1580, and 5000 µg TOS/plate. Stock solutions of the test item at the selected concentrations were confirmed by validated HPLC-PDA analysis for their total protein concentration. Each concentration was plated in triplicate per strain, per condition. Plates were incubated at 37 ± 2 °C for 48 ± 1 h (TA100, TA102) or 72 ± 1 h (TA98, TA1535, TA1537), and revertant colonies were counted manually against historical ranges. Strain-specific positive controls recommended by OECD TG 471 and strain-appropriate solvents were included every run; sterility controls were performed (11, 12). Assay validity required positive controls to yield the expected fold increases and vehicle controls to fall within historical ranges (11). A test item response was considered positive if a biologically relevant, concentration-related increase occurred, typically ≥2-fold for TA98/TA100/TA102 and ≥3-fold for TA1535/TA1537, with reproducibility (11, 12).

### In vitro micronucleus test (OECD TG 487)

Clastogenicity/aneugenicity was evaluated in cultured human peripheral blood lymphocytes under OECD Test No. 487 (9). Cultures from healthy non-smoking male donors were exposed to the test item at 1250, 2500, and 5000 µg TOS/mL for 4 h without S9, 4 h with S9 (followed by recovery), and 24 h without S9. Stock solutions of the test item at the selected concentrations were confirmed by validated HPLC-PDA analysis for their total protein concentration. Cytokinesis-block protocol used cytochalasin B; cytotoxicity was quantified by the cytokinesis-block proliferation index (CBPI) from 1000 cells per culture; micronuclei were scored in 2000 binucleated (BN) cells per concentration (two cultures per condition) using standard morphological criteria (9, 13). Positive controls were vinblastine sulfate (−S9, 4 h), cyclophosphamide (+S9, 4 h), and mitomycin C (−S9, 24 h). Acceptance criteria followed OECD TG 487, including target top-dose cytotoxicity approaching 55 ± 5% where feasible, concurrent positive control responses, and vehicle controls within laboratory historical ranges (9, 13). Per TG 487, if cytotoxicity does not reach approximately 55% despite testing the maximum recommended concentration (5000 µg TOS/mL) with acceptable pH/osmolality and no precipitation, the study remains valid and the top concentration is sufficient for hazard identification.

### In vivo repeated-dose oral toxicity

Animals and husbandry

Sprague Dawley rats for both the 28-day and 90-day studies were sourced from Hylasco Biotechnology (India) Pvt. Ltd. For the 90-day study, males were approximately 4–5 weeks old and females 5–6 weeks old at first dose; Day 1 body-weight ranges were 153.28–191.62 g for males and 145.21–184.69 g for females. For the 28-day study, Sprague Dawley rats of similar age from the same supplier were used; age at first dose and initial body-weight ranges by sex were within the normal range for young, rapidly growing SD rats as reported in the 28-day study report. Animals were acclimated 7 days (males) or 8 days (females). Housing: ≤3 per polycarbonate cage, stainless steel grill tops, autoclaved corncob bedding. Environmental conditions were monitored: 20.2–23.4 °C (28-day study) or 20.8–23.4 °C (90-day study); 46– 68% or 49–68% relative humidity; 12 h light:12 h dark; 10–15 air changes/hour. Diet was gamma-irradiated rodent maintenance feed, ad libitum except during fasting procedures; water was RO-treated, UV-exposed, autoclaved, ad libitum. Husbandry followed the facility SOPs and the Guide for the Care and Use of Laboratory Animals (16).

### Study design and randomization (OECD TG 407 and 408)

Two GLP studies were conducted: a 28-day repeated-dose study (OECD TG 407) and a 90-day subchronic study (OECD TG 408) (10, 14). In each study, animals were assigned to four groups (n = 10/sex/group) by body-weight-stratified randomization with ≤±20% variation from sex-specific means. Groups received vehicle control (0), 500, 1000, or 2000 mg TOS/kg/day at 10 mL/kg by oral gavage once daily for 28 or 90 consecutive days, respectively. Necropsy occurred on Day 29 (28-day) or Day 91 (90-day). Group identities were maintained for dose preparation and administration; pathology evaluation followed GLP procedures.

### Dosing and in-life observations

Dose volumes were adjusted weekly from body weight. Animals were observed twice daily for mortality/morbidity and at least once daily for clinical signs. Detailed clinical examinations were performed pre-dose and weekly. Cage rotation occurred weekly. Body weight was recorded at randomization, Day 1 pre-dose, weekly thereafter, and terminally (fasted). Body-weight gains were computed relative to Day 1. Feed consumption was determined cage-wise at weekly intervals using the formula: (offered − leftover)/(animals per cage × days). Ophthalmology was performed during acclimatization for all the animals and in Week 13 on control and high-dose groups by indirect ophthalmoscopy per SOP. Functional observation battery (FOB) was conducted during Week 4 (28-day: control and high-dose groups) and Week 12 (90-day: control and high-dose groups), including home-cage, handling, open-field, reflex, neuromuscular (grip strength, hind-limb splay), and rectal temperature measures (10, 14).

### Clinical pathology

For terminal clinical pathology (Day 29 or Day 91), animals were fasted overnight with water *ad libitum*. Blood was collected from the retro-orbital plexus under isoflurane. Anticoagulants were 10% K2EDTA (2 mg/mL blood) for hematology, 3.2% sodium citrate (3.2 mg/mL blood) for coagulation, and lithium heparin (20 IU/mL blood) for clinical chemistry. Plasma/serum were obtained by centrifugation at 4000 rpm, 10 min, 4 °C; serum were stored at approximately −80 °C until analysis.

Urinalysis (Week 4 for 28-day; Week 13 for 90-day) was performed on overnight urine collected in metabolic cages with water available *ad libitum*. Physical parameters included volume, color, and clarity. Test-strip chemistry included pH, specific gravity, occult blood, urobilinogen, ketones, protein, bilirubin, glucose, nitrite, and leukocytes (CLINITEK Status+). Microscopy after centrifugation (1500 rpm, 10 min, 4 °C) assessed epithelial cells, erythrocytes, casts, crystals, white blood cells/pus cells, bacteria, and yeast.

Hematology (e.g., RBC, HGB, HCT, MCV, MCH, MCHC, PLT, WBC and differential) and coagulation (prothrombin time, activated partial thromboplastin time) were measured on an ADVIA 2120 analyzer and TCaog analyzer respectively. Clinical chemistry included albumin, total protein, globulin (calculated), urea nitrogen, creatinine, glucose, cholesterol, LDL, electrolytes (Na^+^, K^+^, Cl^-^, Ca, P), bilirubin, and hepatic enzymes (ALT, AST, ALP, GGT). Panels were selected to support interpretation of observed differences reported in Results (cholesterol, calcium, total protein, globulin, urea nitrogen, glucose, LDL, phosphorus; prothrombin time; mean corpuscular hemoglobin concentration).

### Vaginal cytology

Estrous cycle stage was evaluated on Day 90 by vaginal lavage. Smears were staged as proestrus, estrus, metestrus, or diestrus by light microscopy.

### Necropsy, organ weights, and histopathology

All animals were euthanized under terminal anesthesia followed by exsanguination per SOP on Day 29 or Day 91. A complete gross examination was performed. Absolute and relative organ weights were recorded for brain, heart, liver, spleen, kidneys, adrenals, thymus, lungs, testes/epididymides or ovaries/uterus, and thyroids/parathyroids for the 90-day study. Tissues were fixed in neutral-buffered formalin (testes/eyes in modified Davidsons/Davidson fixative), processed to paraffin, sectioned, and stained with hematoxylin and eosin. Histopathology followed OECD TG 408 lists for comprehensive evaluation in control and high-dose groups. Bone marrow smears were reserved but not evaluated given the absence of hematology or lymphoid organ triggers in these studies.

### Quality assurance, GLP, and ethics

All studies were conducted under GLP and facility SOPs (15). Protocols were approved by the Institutional Animal Ethics Committee (IAEC) at Anthem Biosciences Ltd. (Protocol ABD/IAEC/PR/330 24 25 for the 28-day study and ABD/IAEC/PR/332 24 25 for the 90-day study). Archiving followed facility policy for 9 years after study completion. QA oversight audited protocol, raw data, and reports.

### Statistics

Analyses were performed separately by sex. Endpoints were analyzed by one-way ANOVA followed by Dunnett’s test versus concurrent vehicle control when assumptions were met;. Urinalysis semi-quantitative categories were summarized descriptively. Statistical significance was set at p < 0.05 (two-sided). For in vitro assays, trend analyses and fold-change criteria followed OECD TG 471 and TG 487 (9, 11). Sample size (n = 10/sex/group) and the high-dose selection (2000 mg TOS/kg/day) followed OECD TG 407/408 and prior range-finding data (10, 14).

## Results

### Bacterial reverse-mutation assay (OECD 471)

P24 did not induce mutations in any tester strain in either assay format or metabolic condition. In the plate incorporation method up to 5000 µg TOS/plate, mean revertant counts for all strains with and without S9 were comparable to concurrent vehicle controls and mutagenicity factors (MF = treatment/vehicle) remained close to unity, with no concentration-related increase in any strain. The preincubation method gave the same conclusion, with no cytotoxicity and no dose-related increase in revertants in any strain. All positive controls produced the expected ≥2-fold (TA98/TA100/TA102) or ≥3-fold (TA1535/TA1537) increases, confirming assay sensitivity. Stock solutions used for the assay were confirmed by validated HPLC-PDA analysis for total protein concentration and met predefined acceptance criteria (mean recovery 92.95–94.67%, %RSD ≤ 20%). Top-dose (5000 µg TOS/plate) revertant counts and mutagenicity factors for each strain and test format are summarized in Table 1. The test item was therefore considered non-mutagenic under the conditions of OECD 471 (11–12). (Table 1)

**Table 1.**
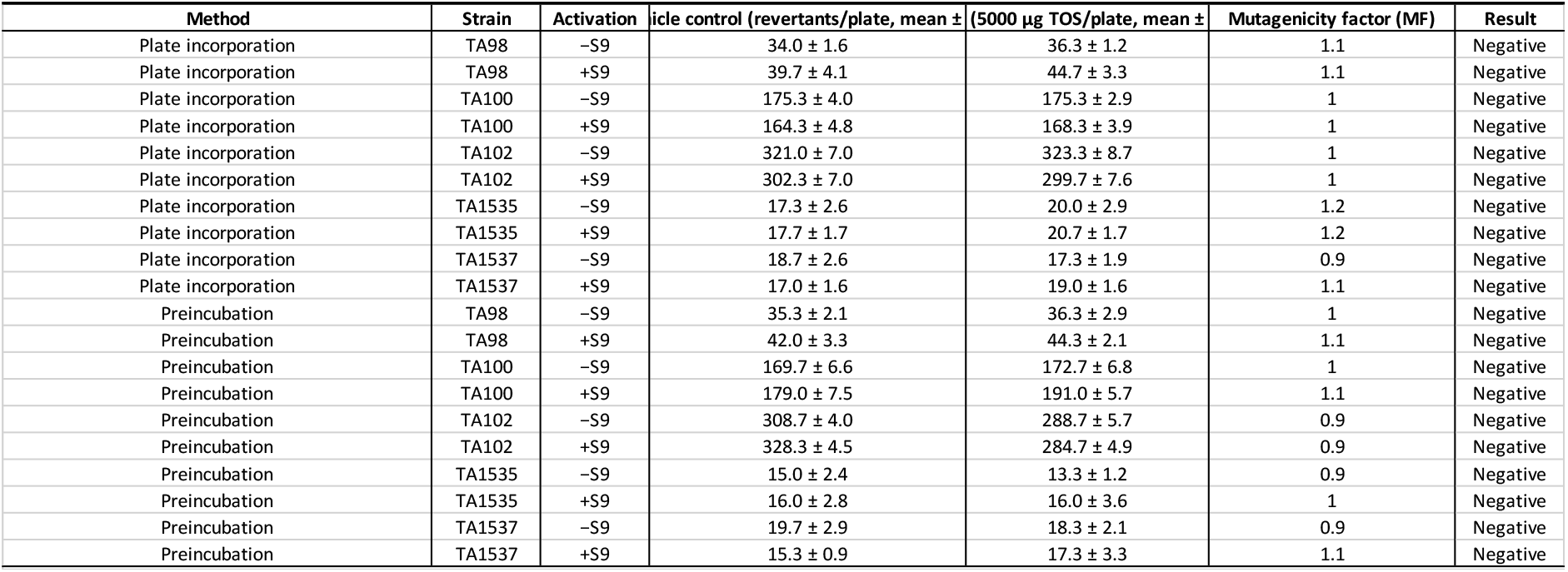
Mean revertant counts (± SD) and mutagenicity factors (MF = treated/vehicle) at the top concentration of 5000 µg TOS/plate in the bacterial reverse-mutation assay with *Salmonella typhimurium* strains TA98, TA100, TA102, TA1535, and TA1537. Data are shown for plate-incorporation and preincubation methods with and without metabolic activation (−S9, +S9). For all strains and conditions, treated cultures showed revertant numbers comparable to concurrent vehicle controls (MF ≈ 1) and were interpreted as negative for mutagenic activity under OECD TG 471.

### In vitro micronucleus assay (OECD 487)

No clastogenic or aneugenic activity was detected in human lymphocytes up to 5000 µg TOS/mL under any exposure condition. Cytotoxicity at the top concentration remained low: 4 h −S9, 19.33%; 4 h +S9, 15.55%; 24 h −S9, 21.56% (all within recommended windows). Because 5000 µg TOS/mL was the maximum feasible concentration and produced ≤21.6% cytotoxicity with acceptable pH/osmolality and no precipitation, this satisfies TG 487 validity even though the ∼55% target was not reached. Micronucleated binucleate (%MNBN) frequencies at 5000 µg TOS/mL were indistinguishable from concurrent vehicles: 4 h −S9, 0.20% vs 0.15%; 4 h +S9, 0.25% vs 0.10%; 24 h −S9, 0.20% vs 0.10%. Lower concentrations (1250 and 2500 µg TOS/mL) yielded %MNBN values between 0.00% and 0.20% across conditions, without dose–response. All positive controls were significant as expected: vinblastine (−S9, 4 h) 1.55%; cyclophosphamide (+S9, 4 h) 1.45%; mitomycin C (−S9, 24 h) 1.70%. HPLC dose confirmation for low/mid/high stocks (12.5/25/50 mg TOS/mL) was 96.33–97.90% (RSD 0.32–1.11%). P24 was therefore negative in OECD 487 (9). (Figure 1)

**Figure 1.**
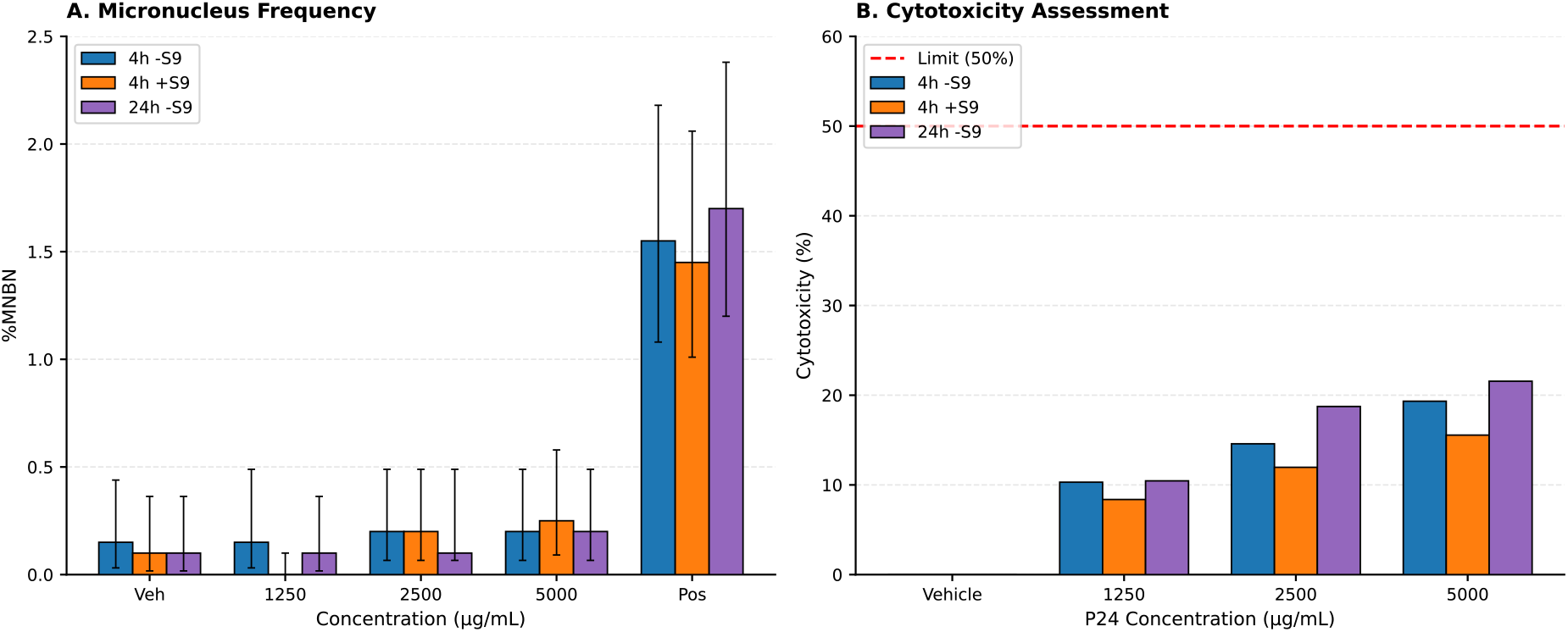
In vitro micronucleus response of human lymphocytes exposed to kumamolisin (P24). Human peripheral blood lymphocytes were treated with P24 at 0, 1250, 2500, and 5000 µg TOS/mL under three exposure conditions: (A) 4 h without metabolic activation (−S9) with recovery, (B) 4 h with metabolic activation (+S9) with recovery, and (C) 24 h without S9. For each condition, the frequency of micronucleated binucleated (%MNBN) cells is plotted as mean ± SD (n = 2 cultures per concentration, 2000 BN cells scored per concentration). (D) Corresponding cytotoxicity, expressed as percent reduction in the cytokinesis-block proliferation index (CBPI) relative to vehicle control, for the same concentrations and exposure conditions. Vinblastine (4 h −S9), cyclophosphamide (4 h +S9), and mitomycin C (24 h −S9) served as positive controls and produced robust increases in %MNBN (not shown). No test-item–related increase in micronuclei or cytotoxicity was observed up to the guideline limit concentration of 5000 µg TOS/mL.

### 28-day repeated-dose oral toxicity study in rats (OECD 407)

In-life observations. No mortality (0/80) or moribundity occurred. Daily clinical signs and weekly detailed clinical examinations were unremarkable in all groups at 0, 500, 1000, and 2000 mg TOS/kg/day. Home-cage, handling, open-field, reflex, neuromuscular, and physiological domains in the functional-observation battery (FOB) were comparable between control and high-dose cohorts, with no treatment-related findings.

Body weight and feed intake. Weekly body weight, body-weight gain, and average weekly feed consumption were unaffected in both sexes at all doses. There were no dose-related trends and no statistically significant deviations that persisted across time points.

Urinalysis. Group means for pH, specific gravity, urobilinogen, bilirubin, ketones, glucose, protein, nitrite, and blood were comparable to controls. High-dose females showed a higher urine volume with small decreases in pH and specific gravity, but values remained within standard Sprague–Dawley ranges and lacked any histopathological correlates in kidney; these changes were considered not adverse.

Clinical pathology. Hematology and coagulation were comparable to controls, with only marginal, single-sex, non–dose-related differences: MCHC +2.2% in males at 500 mg/kg/day and PT −9.7% in females at 2000 mg/kg/day, the latter in the absence of liver histopathology. Clinical chemistry showed small, isolated shifts without internal consistency or organ correlates: cholesterol +18.8% in males at 500 mg/kg/day and calcium −3.1% in females at 500 mg/kg/day; both were judged incidental.

Pathology. Gross necropsy revealed no test-item-related lesions. Statistically significant heart-weight decreases in females at 500 and 1000 mg/kg/day were ≤14% for absolute and relative weights, were not dose-related, occurred in one sex only, and had no microscopic correlates; they were considered non-adverse. Histopathology showed no P24-related findings at 2000 mg/kg/day in either sex. Isolated, minimal changes (e.g., unilateral diffuse mild renal pelvis dilation and chronic pulmonary inflammation in individual high-dose males; minimal focal hepatocellular necrosis and minimal focal bilateral renal infarcts in individual high-dose females) were sporadic background lesions for the strain and age and showed no treatment pattern. The 28-day NOAEL was 2000 mg TOS/kg/day, the highest dose tested (14).

Given the absence of adverse findings at 2000 mg TOS/kg/day over 28 days, the 90-day study was conducted at the same top dose with identical vehicle, dose volume, and analytical controls for formulation homogeneity and stability. The next section reports the subchronic outcomes in detail.

### Subchronic 90-day oral toxicity in rats (OECD 408)

#### Study overview and exposure verification

Sprague Dawley rats (n=80; 10/sex/group) from Hylasco Biotechnology were assigned to four groups and dosed by oral gavage once daily for 90 consecutive days at 0, 500, 1000, or 2000 mg TOS/kg bw/day in ultrapure water (10 mL/kg). At treatment start, males were 4–5 weeks old and females 5–6 weeks old; day-1 body-weight ranges were 153.28–191.62 g (males) and 145.21–184.69 g (females). Animal rooms were maintained at 20.8–23.4 °C and 49–68% RH on a 12:12 h light:dark cycle. Dose-formulation accuracy and homogeneity were confirmed by HPLC: day 0 recoveries were 102.50 ± 0.38% (50 mg TOS/mL; %RSD 0.37), 102.38 ± 1.11% (100 mg/mL; %RSD 1.08), and 97.75 ± 0.81% (200 mg/mL; %RSD 0.83). Day 90 recoveries were 101.62 ± 0.74% (50 mg/mL; %RSD 0.73), 100.95 ± 0.98% (100 mg/mL; %RSD 0.97), and 99.43 ± 0.90% (200 mg/mL; %RSD 0.91). Vehicle blanks were negative. Study conduct followed GLP and OECD TG 408 (10,15). Results are presented in a quantitative narrative consistent with the Foods journal style.

#### In-life observations

No deaths or moribund events occurred. Daily clinical signs and weekly detailed examinations were normal in all animals throughout the 91 observation days. Ophthalmology at week 13 showed no abnormalities in high-dose animals of either sex; lower-dose groups were not examined due to the absence of findings at 2000 mg TOS/kg/day.

#### Body weight and growth

Mean terminal body weights (day 90) in males were 560.71 g (control), 597.85 g (+37.14 g, +6.6% vs control), 567.81 g (+7.10 g, +1.3%), and 537.68 g (−23.03 g, −4.1%) for 0, 500, 1000, and 2000 mg TOS/kg/day, respectively. Females were 328.09 g (control), 341.91 g (+13.82 g, +4.2%), 328.81 g (+0.72 g, +0.2%), and 341.13 g (+13.04 g, +4.0%). Cumulative body-weight gains from day 1 to 90 were 218.95%, 240.86%, 220.14%, and 203.00% in males and 100.55%, 104.99%, 99.25%, and 104.02% in females (0→2000 mg TOS/kg/day). These marginal between-group differences lacked toxicological patterning and were not interpreted as adverse (Figure 2A, 2B).

**Figure 2.**
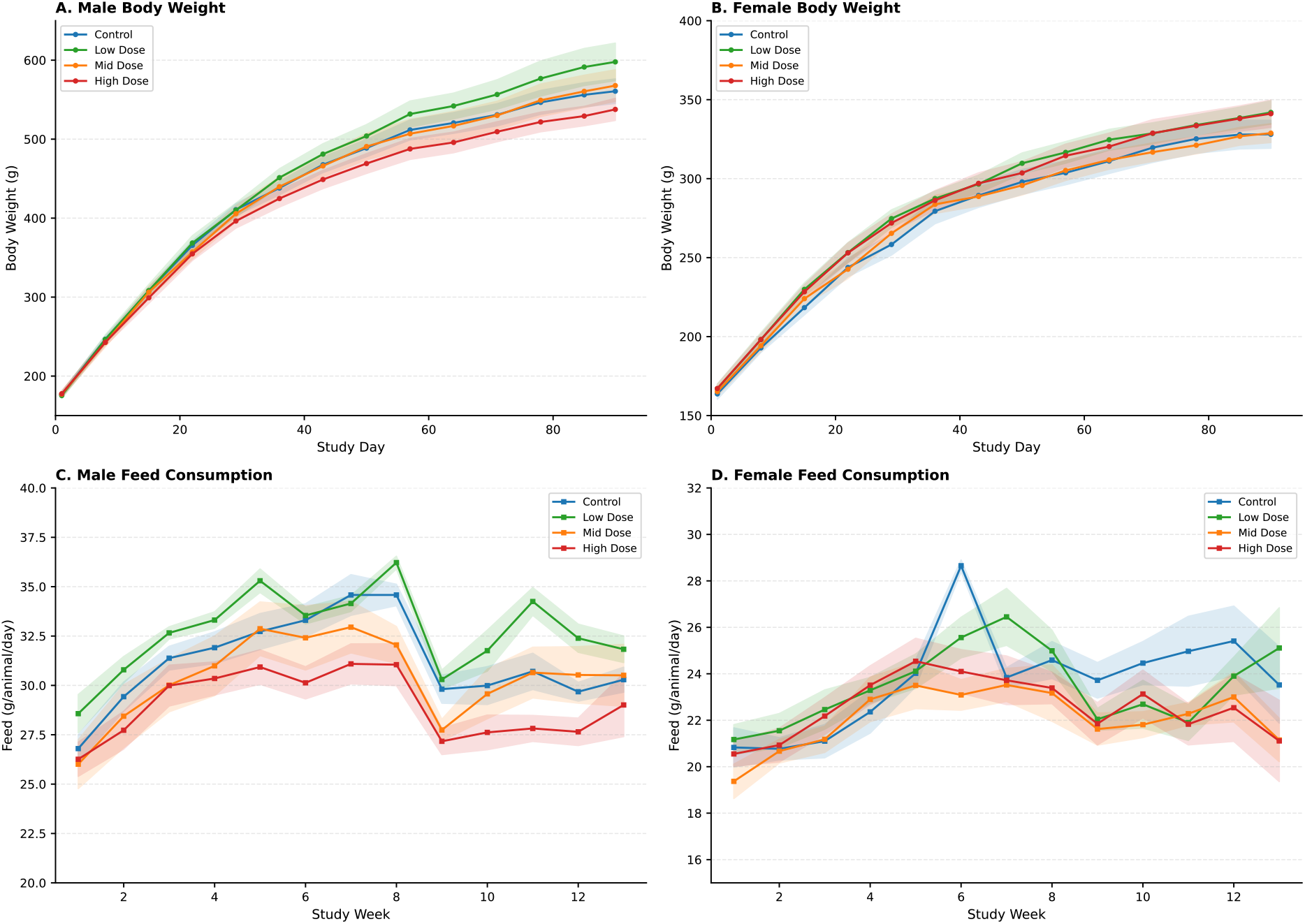
Body weight and feed consumption in Sprague Dawley rats during the 90-day oral toxicity study. Sprague Dawley rats (10/sex/group) received kumamolisin (P24) by oral gavage once daily at 0, 500, 1000, or 2000 mg TOS/kg bw/day for 90 days. (A) Mean body weight in males and (B) females over time (Days 1–90). (C) Mean weekly feed consumption in males and (D) females on a per-animal basis. Values are mean ± SD. Small between-group differences in body weight gain and feed intake showed no consistent dose-response pattern and were considered not adverse. Feed consumption. Cage-wise weekly means showed isolated differences without dose–response. In males: week 11 low-dose was higher than control by +11.5% (34.25 vs 30.71 g/animal/day); weeks 8 and 9 high-dose was lower than control by −10.2% (31.05 vs 34.58) and −8.8% (27.17 vs 29.81), respectively. In females at week 6: low-, mid-, and high-dose groups were −10.8% (25.56 vs 28.65), −19.4% (23.09 vs 28.65), and −15.9% (24.10 vs 28.65) vs control. Given inconsistency across weeks and sexes, and the absence of concordant clinical or anatomical findings, these shifts were considered non-adverse (Figure 2C, 2D).

#### Functional observation battery (week 13)

Qualitative measures (posture, respiration, gait, mobility, reflexes) were normal in all animals tested (G1 vs G4). Quantitative endpoints showed close concordance between control and high-dose: urinations (males 2.00 vs 2.10; females 0.50 vs 0.50), defecations (males 1.00 vs 1.00; females 0.00 vs 0.00), rearing (males 6.70 vs 6.10; females 9.30 vs 9.90), rectal temperature (°C; males 37.29 vs 37.36; females 37.00 vs 37.09), and landing foot splay (cm; males 6.91 vs 6.97; females 6.58 vs 6.81). No neurofunctional signals of systemic toxicity were identified (16).

#### Urinalysis (week 13)

Male mean pH was 7.30–7.50 with specific gravity 1.015–1.016 across groups; female mean pH was 7.00–7.25 with specific gravity 1.015–1.017. Semi-quantitative chemistries (e.g., protein, glucose, ketone bodies, bilirubin, blood) and microscopy (cells, casts, crystals, bacteria, yeast) were comparable to controls for both sexes. No item-related renal signal was detected by urinalysis.

#### Clinical chemistry

Group means were generally aligned with controls. Statistically significant differences were limited to males for total protein (+5.8% mid, 7.30 g/dL; +9.1% high, 7.53 g/dL; control 6.90), globulin (+9.3% high, 6.20 vs 5.67 g/dL), blood urea nitrogen (+16.2% mid, 13.60 mg/dL; +20.5% high, 14.10 mg/dL; control 11.70), glucose (+11.9% mid, 118.50 mg/dL; +10.7% high, 117.20 mg/dL; control 105.90), LDL cholesterol (+84.6% mid, 14.40 mg/dL; +110.3% high, 16.40 mg/dL; control 7.80), and calcium (−5.3% mid, 9.27 mg/dL; −6.3% high, 9.17 mg/dL; control 9.79) (Figure 3A). Although the LDL percent changes appear large, the absolute LDL values remained very low (14–16 mg/dL), which is typical for Sprague Dawley rats, whose lipid profiles are HDL-dominant and show broad physiological variability at these concentrations. The shifts were single-sex, modest in absolute terms, within historical control ranges, and occurred without associated changes in total cholesterol, triglycerides, or liver histopathology, indicating they were not toxicologically meaningful. In females, phosphorus was higher in mid and high groups (+17.2%; 5.38 vs 4.59 mg/dL), but values were within historical variability and lacked supporting clinical or anatomical findings (Figure 3B). Overall, all noted differences lacked dose-coherent progression across sexes, had no macroscopic or microscopic correlates, and were therefore interpreted as incidental and non-adverse.

**Figure 3.**
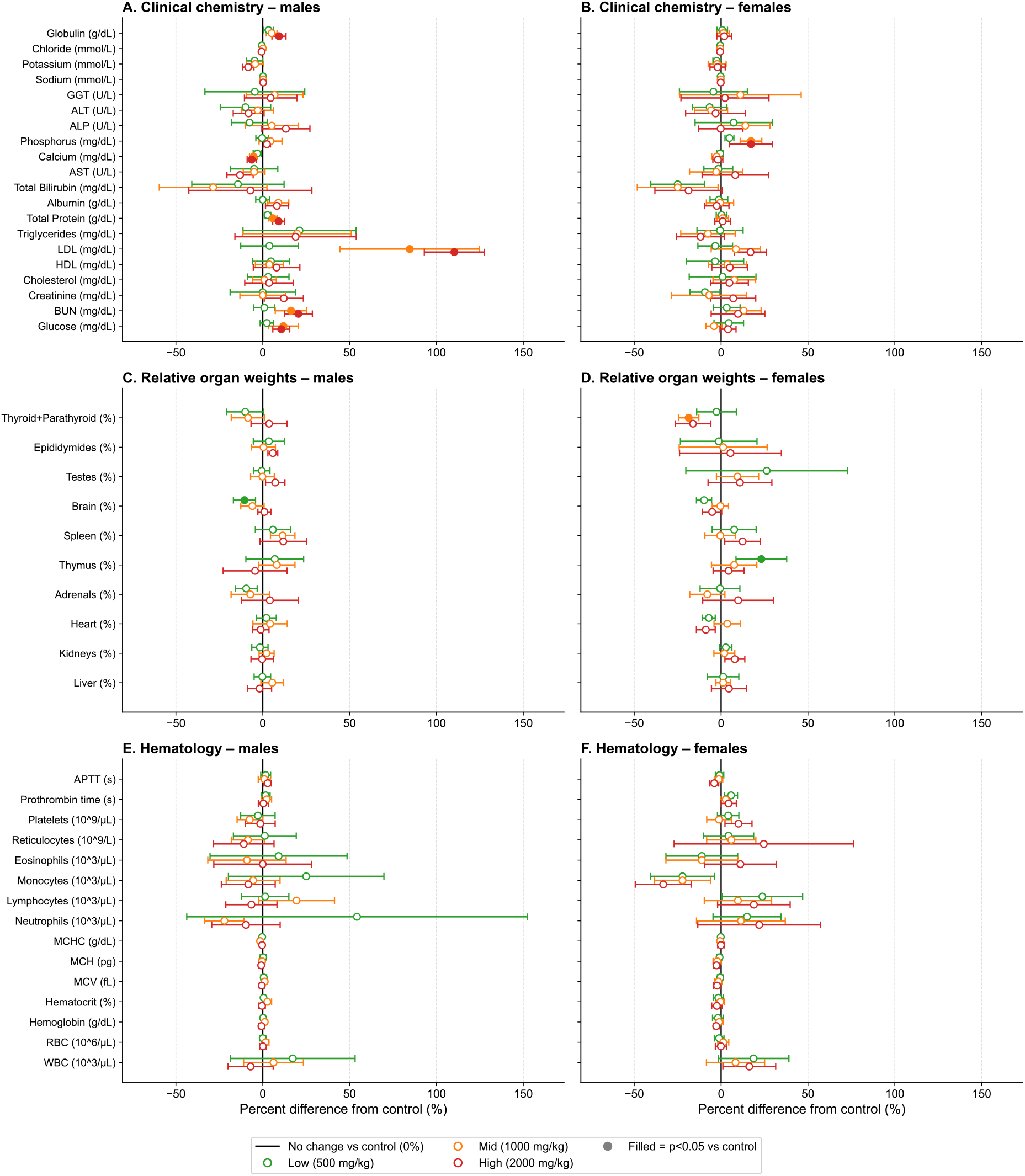
Clinical chemistry, relative organ weights, and hematology in Sprague Dawley rats after 90 days of oral exposure to kumamolisin (P24). Forest-plot summary of percent differences from concurrent control (0 mg TOS/kg bw/day) with 95% confidence intervals for selected endpoints in males and females (10/sex/group) at 500, 1000, and 2000 mg TOS/kg bw/day. (A) Male clinical chemistry parameters and (B) female clinical chemistry parameters, including total protein, globulin, blood urea nitrogen, glucose, LDL cholesterol, calcium, and phosphorus. (C) Male relative organ weights and (D) female relative organ weights (organ weight as % of terminal body weight) for liver, kidneys, heart, adrenals, thymus, spleen, brain, and reproductive organs. (E) Male hematology indices and (F) female hematology indices, including red and white blood cell counts, hemoglobin, hematocrit, red cell indices, platelets, and coagulation times. Each point represents the mean percent difference from control; error bars show the 95% CI. Filled symbols denote statistically significant differences versus control (p < 0.05). All changes were small in magnitude, lacked coherent dose-response or cross-sex concordance, and had no supporting macroscopic or microscopic findings, and were therefore interpreted as incidental and non-adverse.

#### Hematology and coagulation

No statistically significant between-group differences were detected. Illustrative ranges encompassed WBC 8.40–10.59×10^3/µL (males) and 5.30– 6.29×10^3/µL (females), and prothrombin time 26.19–26.79 s (males) and 22.84–24.15 s (females). The absence of shifts in erythroid indices or platelet parameters further supported a lack of hematologic toxicity. These findings are summarized in Figure 3E and 3F.

#### Estrous cyclicity (day 90)

Vaginal cytology distributions were typical for Sprague Dawley rats with no treatment-related disturbance: control and high-dose groups each had 2 proestrus, 4 estrus, 1 metestrus, and 3 diestrus animals; low- and mid-dose distributions were similarly balanced (2/3/2/3 and 4/2/1/3 across stages, respectively).

#### Gross pathology and organ weights

Terminal necropsy revealed no external or internal gross lesions in any animal (NAD, 40/40 males; 40/40 females). Absolute and relative organ weights were comparable to controls with a few statistically significant differences lacking dose dependence or microscopic correlates. Examples include absolute kidney weight in high-dose females 2.45 g vs control 2.16 g (+0.314 g; +14.5%) and absolute thymus weight in low-dose females 0.505 g vs 0.385 g (+0.120 g; +31.2%). Relative weights showed isolated shifts: male brain 0.401% vs 0.448% (low vs control; −0.047 percentage points; −10.5%), female thymus 0.156% vs 0.127% (low vs control; +0.0295 pp; +23.2%), and female thyroid + parathyroid 0.00583% vs 0.00717% (mid vs control; −0.00134 pp; −18.7%). Given the single-sex occurrence, lack of monotonicity, small effect sizes, and normal histology, these differences were considered non-adverse (Figure 3C, 3D).

#### Histopathology

Comprehensive microscopic evaluation detected no test-item–related changes in any tissue at 2000 mg TOS/kg/day in either sex. Bone marrow smears were not evaluated due to the absence of hematologic signals. The overall pathology profile did not indicate target-organ toxicity.

#### NOAEL

Under the conditions of this OECD 408 study, the no-observed-adverse-effect level for Kumamolisin (P24) was 2000 mg TOS/kg bw/day, the highest dose tested (16).

## Conclusion

A standardized toxicology battery found no evidence that the Bacillus sp. MN-32 kumamolisin exhibits genotoxic or systemic toxic effects under guideline conditions relevant to oral exposure from food uses. In two in vitro assays, the enzyme was negative for mutagenicity in Salmonella typhimurium (TA98, TA100, TA102, TA1535, TA1537) up to 5000 µg TOS/plate across plate-incorporation and preincubation formats with and without metabolic activation, with no cytotoxicity at any concentration tested; dose verifications were within 92.95–94.67% recovery (RSD ≤ 20%). In cultured human peripheral blood lymphocytes, kumamolisin produced no clastogenic or aneugenic response at 1250, 2500, or 5000 µg TOS/mL under 4-h (±S9) and 24-h (−S9) exposures; cytotoxicity at the top dose was ≤ 21.56% and stock-solution recoveries were 96.33–97.90% (RSD ≤ 1.11%). Positive controls performed as expected in both test systems, confirming assay sensitivity.

Oral repeat-dose studies in Sprague Dawley rats identified no target-organ toxicity through 2000 mg TOS/kg bw/day, the limit dose for food enzymes. In the 28-day study (n = 10/sex/group; 0, 500, 1000, 2000 mg TOS/kg/day), there were no test-item–related effects on clinical signs, mortality, weekly body weight, feed consumption, functional observations, urinalysis, hematology, plasma chemistry, serum hormonal, organ weights, or histopathology; the NOAEL was 2000 mg TOS/kg/day. The pivotal 90-day study (n = 10/sex/group; same dose design) likewise showed 0/10 mortality in all groups and no treatment-related findings in clinical observations, ophthalmology (week 13, NAD in 10/10/sex at 2000 mg/kg/day), or functional observation battery. Mean terminal body weights and body-weight gains were comparable to controls (e.g., day-1→90 gain, males: 203.00–240.86%; females: 99.25–104.99% across groups). Group-wise, statistically significant differences in select chemistry endpoints. Males: total protein +5.8% (G3) and +9.1% (G4), globulin +9.3% (G4), BUN +16.2% (G3) and +20.5% (G4), glucose +11.9% (G3) and +10.7% (G4), LDL +84.6% (G3) and +110.3% (G4); calcium −5.3% (G3) and −6.3% (G4); females: phosphorus +17.2% (G3, G4)—remained within historical control ranges, lacked dose-consistent progression in both sexes, and had no macroscopic or microscopic correlates. Organ-weight differences that reached statistical significance were marginal, sporadic, single-sex, and non–dose-responsive (e.g., kidney, thymus); comprehensive histopathology at 2000 mg/kg/day revealed no test-item–related lesions. The subchronic NOAEL was therefore 2000 mg TOS/kg bw/day.

Quality controls across studies support the robustness of these conclusions. Clinical batches were analyzed at preparation for concentration, homogeneity, and stability: recoveries were 96–103% at 50–200 mg TOS/mL with %RSD ≤ 1.2%; preparations were stable for 7 h at 22 ± 3 °C and 72 h at 2–8 °C. Specific pathogen free (SPF) animals were sourced healthy, randomized by body weight (≤ ±20% of sex-specific means), and maintained under monitored environmental conditions; sample sizes (10/sex/group) met OECD power expectations for subchronic hazard identification.

For risk characterization, the 90-day NOAEL (2000 mg TOS/kg bw/day) compared to a conservative 90th-percentile intake estimate (12 mg TOS/kg bw/day for young children) yields a margin of safety of 167-fold. This exceeds the default 100-fold composite uncertainty factor conventionally applied to subchronic rodent NOAELs for human dietary exposure, indicating that the tested kumamolisin presents a low toxicological concern at anticipated use levels.

Limitations inherent to the dataset include reliance on a single species and strain, one enzyme batch, and the absence of bone-marrow smears (not indicated by hematology or lymphoid histology). These do not materially affect the weight of evidence given the concordant negative genotoxicity, absence of adverse findings at the 2000 mg/kg/day limit dose over 90 days, and large human exposure margin. Taken together, the data demonstrate that kumamolisin from Bacillus sp. MN-32 is non-genotoxic in vitro and lacks systemic toxicity in rats up to 2000 mg TOS/kg bw/day, supporting a favorable safety profile for oral exposure under intended food-use scenarios.

